# Environmental dipolar relaxation during excited state proton transfer in Green Fluorescent Protein

**DOI:** 10.1101/2025.11.20.688880

**Authors:** Moona Kurttila, Inês S. Camacho, Athena Zitti, James A. Platts, Javier Garcia-Ruiz, Richard W. Clarke, Christopher R. Pudney, D. Dafydd Jones, Alex R. Jones

## Abstract

Engineered variants of Green Fluorescent Protein (GFP) are in widespread use as genetically-encoded labels for biological imaging. Excitation of its neutral phenolic ground state chromophore populates an emissive anionic phenolate state *via* excited state proton transfer (ESPT) to E222 along a molecular wire. Computational studies indicate that ESPT fits to an electronically adiabatic rate expression. Although reactions that operate close to the adiabatic limit can be sensitive to environmental dipolar relaxation, little is known about the role of such relaxation during ESPT in GFP. Here, we present the first experimental evidence that dipolar relaxation of the protein matrix occurs during ESPT in response to the change in electron density distribution in the excited state GFP chromophore and is a key determinant of the reaction pathway. Using fluorescence spectroscopy, we excited along the red edge of the neutral phenolic ground state absorption band of several GFP variants with differing chromophore environments. Instead of resulting in a significant red shift of the centre of spectral mass (CSM) of the emission spectra common for biological chromophores such as tryptophan, the CSM of each GFP variant remains almost unchanged as a function of excitation wavelength. This is consistent with each sub-population that is selectively excited along the red edge reaching the same, fully relaxed state before emission of a photon. The kinetics of environmental dipolar relaxation are therefore on the same sub-nanosecond timescale as ESPT, which provides an explanation for its adiabatic nature and could inform the rational design of novel fluorescent proteins with tailored photophysics.

## Introduction

Molecular chromophores are found throughout nature. Their varied, conjugated structures mean they absorb wavelengths from the ultraviolet B (UVB, ∼260 nm)^1–3^ across the visible to the near infrared (NIR, ∼ 1000 nm).^4^ Some, like the Green Fluorescent Protein (GFP) chromophore, have evolved (and been further engineered) to be efficient re-emitters of light.^5–6^ Others use the absorbed energy to trigger chemical or structural changes,^7–10^ but in many contexts exploit their molecular properties to support light-independent redox^11^ or structural^12^ functions. Irrespective of their primary function, light absorption results in a change in electron density distribution within the chromophore, and hence a change in the magnitude and direction of its dipole moment. Such a change will in turn elicit some sort of response from the environment, such as dipolar relaxation of neighbouring molecules (solvent, protein side chains, etc.) to accommodate the changes in dipole moment.^13^ If the chromophore is fluorescent, this environmental response can influence the spectral properties of the emitted light. In particular, if excitation occurs at increasingly longer wavelengths along the red edge of the chromophore’s absorption band it is possible to observe a shift in the inhomogeneous broadening of the resulting emission spectra. Careful analysis of such ‘red edge excitation shift’ (REES) effects can inform on the environmental dynamics of the chromophore,^14–15^ including in biological systems. ^16^ In this study, we use REES to investigate the environmental dynamics of several GFP variants. We find a near-complete absence of a red edge shift, in stark contrast to typical biological chromophores. These results provide direct experimental evidence that the protein matrix undergoes sub-nanosecond dipolar relaxation that is concomitant with excited-state proton transfer (ESPT), thereby explaining the reaction’s adiabatic nature and revealing environmental dynamics as a critical component of the proton transfer pathway.

## Results

To illustrate the REES effect, we have probed the environmental dynamics following the photoexcitation of the amino acid tryptophan (Trp) in water^17^ using fluorescence spectroscopy (Figure 1, see Supporting Information for methods). The nature of the conjugated indole moiety of the Trp sidechain has three main implications in context: i) it is an intrinsic fluorescent reporter in proteins, absorbing UVB (*λ*_max_ ∼280 nm) light and emitting in the UVA (*λ*_max_ ∼350 nm); ii) the steric bulk of its sidechain compared to other canonical amino acids coupled with its ability to contribute to both hydrophobic and H-bond interactions means it often plays a structural role; and iii) it can stabilize charge and unpaired electrons and is thus redox-active. Its fluorescence signal can therefore report on both structural and functional changes in proteins. Following photoexcitation at its absorbance maximum in water (*hυ*_FC_, Figure 1, centre), although the direction and magnitude of the Trp dipole moment changes (Figure S1a and Table S1), the solvent dipoles are initially arranged around the resulting Frank-Condon (FC) state in a way that is still complementary to the ground state dipole moment of Trp. Following vibrational relaxation (*k*_v_), depending on the relative kinetics, the Trp might fluoresce from the FC state (*k*_F1_, *hυ*_1_) or the solvent dipoles might relax (*k*_R_, dipolar relaxation) to complement the excited state dipole before fluorescence (*k*_F2_, *hυ*_2_) from the ‘relaxed’ state (R state).

**Figure 1.**
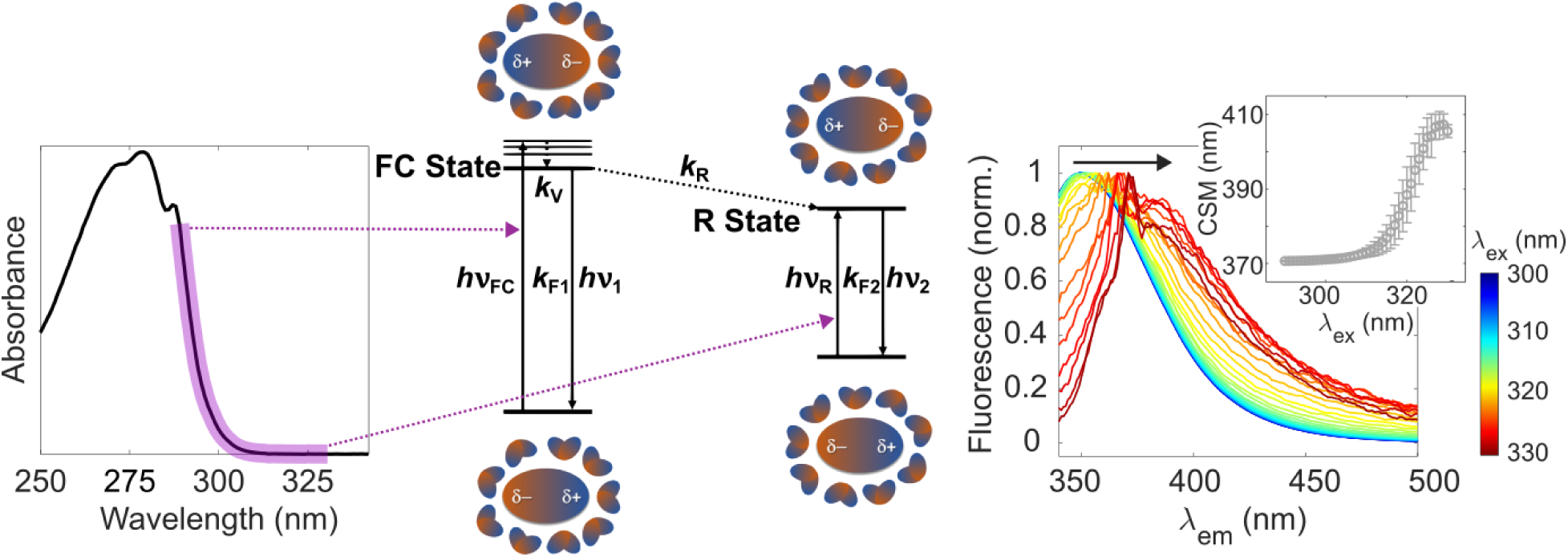
Red-edge excitation shift (REES) effect illustrated using the example of Trp in water. Exciting along the red edge of the Trp absorption spectrum (left) selects for different populations between the high and low limits of the range of interaction energies between the chromophore and solvent dipoles (centre). This results in a change of the inhomogeneous broadening of the Trp emission spectra (right), which can be analysed by plotting the centre of spectral mass (CSM) as a function of excitation wavelength (*λ*_ex_, right inset). See main text for full description.

In the Trp ground state, there are various subpopulations where the solvent dipoles are arranged between the limit where they are entirely complementary to the ground state dipole (main population) and the limit where they are entirely complementary to the excited state dipole (minor population). Exciting Trp along the red edge of its absorption spectrum in a stepwise manner (Figure 1, left) selects for each of these sub-populations. If *k*_F_ ≥*k*_R_, excitation close to the absorbance maximum results in Trp mostly emitting from the FC state, whereas excitation at the longest wavelength of the red edge only populates the R state (*hυ*_R_), and emission is thus only from this state. The peak position and extent of inhomogeneous broadening in the emission spectra change accordingly in a manner that is independent of the emission intensity (Figure 1, right). By integrating the area under each emission spectrum, the centre of spectral mass (CSM, see equation 1 in the Supporting Information) can be determined and plot as a function of excitation wavelength (*λ*_ex_), yielding a sigmoidal relationship (Figure 1, right inset). The nature of the sigmoidal relationship varies depending on the environmental dipolar relaxation around the chromophore. Thus, spectroscopic access to the ensemble of interaction energies between the Trp chromophore and its environment is available.^15^ This fact has been exploited to provide insights into the structure of Trp-containing proteins and peptides using REES, including their conformational dynamics and stability.e.g., ^16, 18–22^

We have used REES to elucidate the nature of the environmental response to excitation of the neutral phenolic GFP ground state chromophore and conversion to the emissive anionic phenolate state *via* ESPT.^23–25^ In *Aequorea victoria* GFP (wildtype, *Av*GFP), the chromophore (CRO) is formed by the covalent rearrangement of three contiguous residues (S65-Y66-G67, Figure 2a).^26^ The chromophore lies within a helical segment that transverses the centre of the β-barrel structure.^27^ The phenol hydroxyl group of the chromophore exists as two states in *Av*GFP: the major, neutral phenolic form (CRO-A state) and a minor, phenolate anionic form (CRO-B state; Figures 2b and c).^28–29^ Mutations^30–32^ and environmental (e.g. pH^33^) changes can change the relative populations of the two states. The chromophore has two distinctive deprotonated emissive excited states, the B* and the I* states. The B* state can be populated directly by excitation of the CRO-B state (*λ*_max_ ∼ 475 nm) and emits at *λ*_max_ ∼ 503 nm.^26, 34^ Alternatively, an intermediate emissive state (I* state) can be populated indirectly by excitation of the CRO-A state (*λ*_max_ ∼ 395 nm), which triggers a sub-nanosecond ESPT from the chromophore (A* state) to E222 *via* a molecular wire comprising a water molecule (W22) and S205 (Figure 2a).^35–36^ This pathway results in slightly red-shifted emission (*λ*_max_ ∼ 508 nm) in comparison to emission from B* state. The structural environment of the I* state remains similar to the A state; the structure can relax to the B* state (Figure 2b)^37^ but the barrier to this is relatively high and is thought to be rare on the fluorescence lifetime.^38^ GFP has been extensively engineered for this fluorescence to aid with imaging within cells, but what can the same fluorescence tell us about the dipolar dynamics around the chromophore?

**Figure 2.**
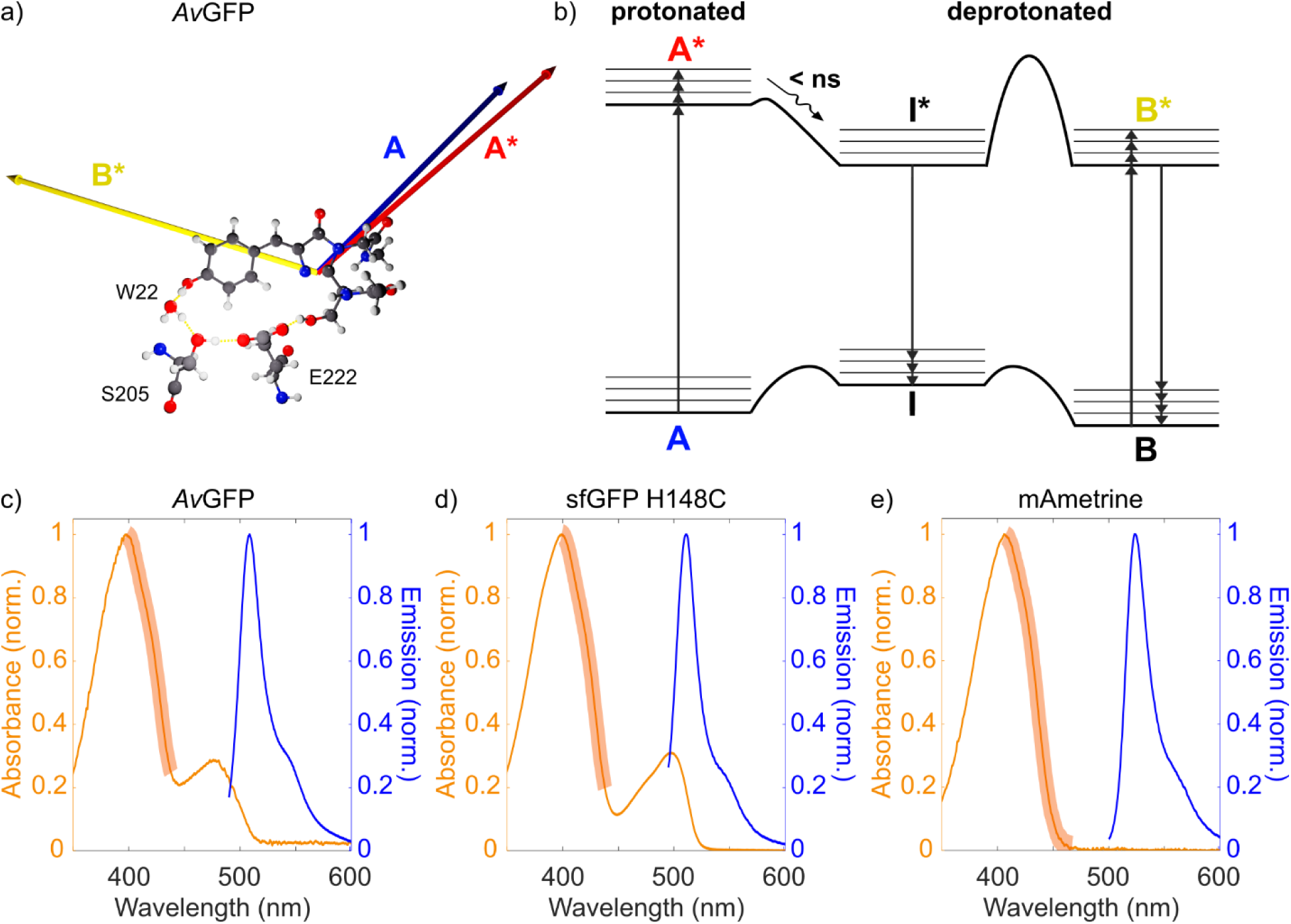
**a)** GFP chromophore and its dipole moments for *Av*GFP. The water (W22) and residues (S205, E222) involved in the ESPT ‘wire’ are highlighted. The directions and relative amplitudes (arrow length) of the dipole moments have been calculated for the chromophore (Table S1) and are indicated for neutral ground (A, red) and excited (A*, blue) states, and the anionic excited state (B*, yellow). **b)** Jablonski diagram showing direct (B→B*) and indirect (A→A*→I*) population of the fluorescent, anionic excited state. Adapted from reference^36^ where relative energy levels are determined for *Av*GFP and could vary between GFP variants; colour coding the same as panel (a). **c-e)** Absorption (orange) and emission (blue) spectra of the three GFP variants measured: *Av*GFP (**c**), sfGFP H148C (**d**), and mAmetrine (**e**). The range of REES excitation wavelengths are highlighted (light orange).

Here, we have focused on *Av*GFP and two variants engineered from *Av*GFP (Figure S1c). The sfGFP H148C variant is an engineered version of the “superfolder” GFP^39^ with the H148C mutation to promote formation of CRO-OH.^39–40^ The spectral properties of sfGFP H148C confirm that the CRO-A (*λ*_max_ 398 nm) dominates over the CRO-B (*λ*_max_ 497 nm; Figure 2d). The second variant, mAmetrine,^41^ is also an extensively engineered version of *Av*GFP (Figure S1c) that almost exclusively populates the CRO-A ground state (single *λ*_max_ at 406 nm) and has a very large Stokes shift owing to a chromophore environment that includes π-π stacking interactions with the chromophore (Figure 2e). By acquiring REES data for GFP variants with a dominant A band in its absorption spectrum and a large Stokes shift (Figures 2c-e), we maximise the extent of the red edge we access before the excitation light interferes with the emission signal and hence maximise the coverage of environmental sub-populations of the CRO-A state. This enabled us to make inferences about the environmental dipolar relaxation dynamics during ESPT.

In sharp contrast to Trp in water (Figure 1), an almost negligible REES effect is observed for each of the GFP variants measured (Figure 3). For *Av*GFP, there is a < 1 nm red shift in the emission profile measured at two different temperatures (20 and 50 °C; Figure 3a), with the observed REES effect at each temperature smaller still (Figure 3a inset). A similar trend is apparent for the *Av*GFP CSM, which shows an essentially flat profile as a function of *λ*_ex_ when plotted on the same scale as the Trp data (Figure 3b). REES effects are expected to reduce at higher temperatures because the rate of dipolar relaxation (*k*_R_) will be greater, and hence emission from the R state is likely to dominate.^42^ Aside from the slight red shift in emission wavelength noted above, any changes in the REES profile with temperature are negligible (Figure S2a), which suggests that emission is predominantly from the R state at both temperatures. At both temperatures, the CSM ultimately *blue* shifts slightly (∼0.5 nm, Figure S2) at the far red edge,^36^ when signal from the B state begins to contribute to the emission signal (Figure S3). To investigate the origin of the flat REES profile, we calculated the dipole moments of the *Av*GFP chromophore (Figures 2a and Table S1; see Supporting Information for methods). Although there is only a small change in magnitude and direction between the A and A* states, consistent with what has been measured experimentally,^43^ there is a significant change in direction between the protonated and excited anionic states. (The dipole moment for the CRO anionic excited state was calculated for B*. Although the protonated E222 near I* might partially influence the dipole, we believe that the values calculated for B* serve as a reasonable approximation.) In principle, therefore, significant relaxation of the environmental dipoles is possible within the emissive state on its fluorescence lifetime (3.1 ns for *Av*GFP,^44^ which is similar to that of Trp, ∼ 2.8 ns^45^).

**Figure 3.**
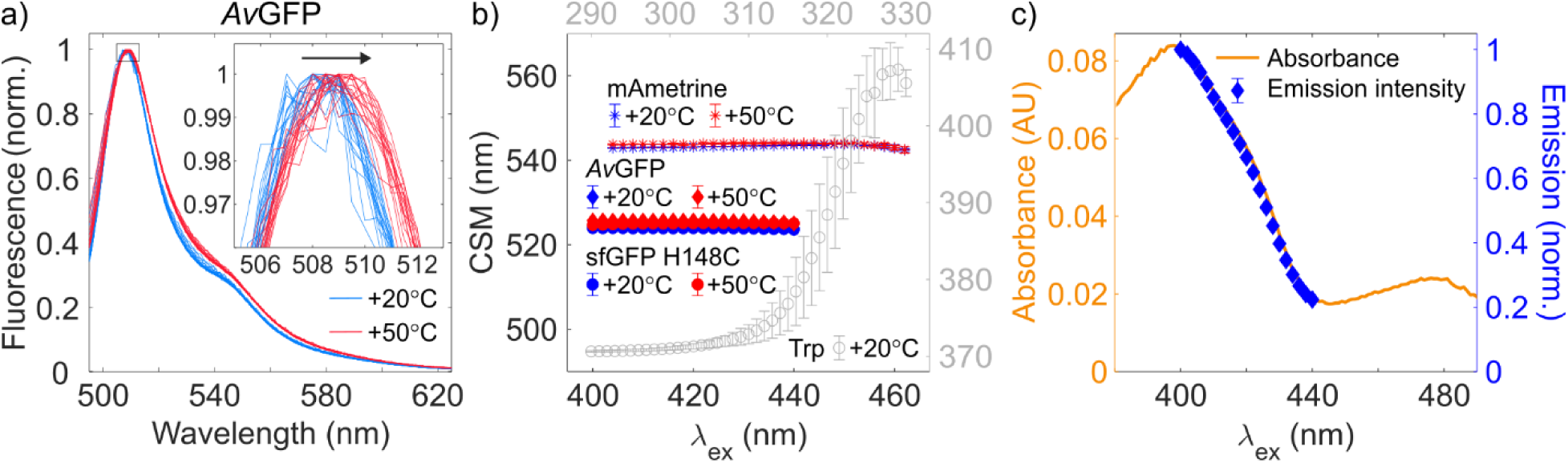
**a)** Normalised emission spectra from *Av*GFP acquired in aqueous solution following excitation along the red edge of its A band (400-432 nm) at 20 °C (blue) and 50 °C (red). A small red shift in the emission peak is evident with the increase in temperature (see also inset). **b)** Average CSM plotted as a function of excitation wavelength (*λ*_ex_) for *Av*GFP, sfGFP-H148C, and mAmetrine at 20 °C (blue) and 50 °C (red); see in-panel legend for data labels; error bars represent one standard deviation. The GFP CSM data (black axes) are plot on the same scale as the equivalent data for Trp (grey data and axes) for comparison. **c)** Red edge of the absorption spectrum of *Av*GFP (orange), overlaid with the average integrated emission intensity of the emission spectrum at each corresponding excitation wavelength (blue) normalized to the emission integral of the series at *λ*_ex_ = 400 nm.

Relatively extensive mutation of the amino acids around the GFP chromophore (Figure S1c), despite significant impact on the absorption spectrum (Figures 2c-e), has little impact on the REES profile (Figures 3b, S2b-c and S4). With regards to sfGFP H148C, the change in the chromophore environment appears to impact the excited state dipole moment, with a larger change in direction of the dipole moment between the A and A* states in sfGFP H148C compared to *Av*GFP and a change of similar magnitude to *Av*GFP between the protonated and deprotonated excited states (Figure S1b and Table S1). Despite this, the REES data show a similarly flat profile until a blue shift at the far-red edge, which overlaps with the CRO-B state excitation (Figures 3b and S2b). The same is true of mAmetrine. The large Stokes shift means we can excite mAmetrine further along the red edge of the absorption spectrum compared to *Av*GFP. Interestingly, we still observe a blue shift in the CSM at the far-red edge despite there being no evidence of an absorption band for the B state, which suggests fluorescence is sensitive to the residual B state population; otherwise, the REES profile is flat (Figures 3b and S2c). In all GFP datasets, the small (∼1 nm) spectral shift is larger between the two temperatures than due to red-edge excitation (Fig. 3b and S3), which emphasises how small the REES effects are for GFP when the CRO-A is excited.

Photoselection of the emissive I* state does not change in GFP as excitation progresses along the red edge of the CRO-A band. A REES study has been conducted previously for the anionic chromophore of enhanced green fluorescent protein (EGFP),^46^ an engineered variant of *Av*GFP with a dominant B absorption band and small Stokes shift.^47–48^ Scanning only ∼25 nm along the red edge of the EGFP B band results in a REES that is significantly larger (∼4 nm) than the < 1 nm REES we observe here scanning ≥40 nm along the red edge of the A bands of *Av*GFP, sfGFP-H148C, and mAmetrine (Figure 3b and S2). This suggests that, when populating the fluorescent, anionic excited state of the chromophore directly (B→B*, Figure 2b), a small but significant range of sub-populations of the environmental dipoles are accessible in the ground state even when exciting only partially along the red edge. If it were practical to do so, we would expect a similar REES effect for the variants measured here. By contrast, exciting along the red edge of the A band populates the fluorescent, anionic excited state indirectly (A→A*→I*, Figure 2b) *via* ESPT. Significant REES effects have been observed previously for emissive states that are populated following excited state reactions, including electron,^49–50^ energy^50^ and proton^51^ transfer, including in proteins.^51^ For example, photoselection of polar, emissive charge-transfer (CT) states at the red edge is possible under conditions where the CT / environment dipolar interactions are equivalent to the R state. This has previously manifested as enhanced population at the red edge of emissive CT states (measured as an increase of emission intensity)^49^ or suppression of emissive states that are in competition with a photoselected CT state.^51^ We have seen already that inhomogeneous broadening of the emission spectra changes little along the red edge of the CRO-A band for several GFP variants (Figures 3a-b, S2, and S4), but what of the emission quantum yield? To quantify the emission intensity and hence the relative yield of the I* state, we integrated under the raw emission spectra at each excitation wavelength. To account for variations in the baseline, each spectrum of the triplicate series at each *λ*_ex_ was normalized to the maximal integral at *λ*_ex_ = 400 nm and then averaged. Comparing this yield at a given wavelength to that expected from the absorption efficiency at the same excitation wavelength (Figure 3c) reveals that, unlike the literature examples discussed above, there is no change in photoselection of the emissive I* state of GFP when populating it *via* ESPT along the red edge of the A band.

## Discussion

The data presented herein are consistent with environmental dipolar relaxation occurring during ESPT (Figure 4). The most likely explanation for why neither any REES effect nor photoselection of the I* state are observed for any GFP variant tested when exciting along the red edge of the A state is that the dipole moments of the residues in the chromophore environment have fully relaxed around the I* dipole by the time a photon is emitted, irrespective of which environmental sub-population of the A ground state is selected upon excitation.^15^ In other words, *k*_R_ > *k*_F_ (Figure 4) and all emission occurs from the dipolar relaxed, R, state of I*. Reactions where charges are transferred can be very sensitive to environmental dipolar effects if they operate close to the adiabatic limit, where there is strong interaction between reactant and product states on a common potential energy surface. Indeed, under such conditions reaction rates can be directly proportional to dipolar relaxation rates in the environment.^52^ In GFP, ESPT along the proton wire from the chromophore to E222 occurs on the picosecond timescale^6, 53^ and fits to an electronically adiabatic rate expression,^6, 54–56^ which suggests *k*_R_ ∼ *k*_ESPT_. Rather than by the intrinsic barrier to proton motion, therefore, the rate of ESPT in GFP appears to be governed by the ability of the surrounding dipolar environment to reorganize in response to the altered charge distribution of the excited state intermediates of the chromophore, as previously discussed for electron transfer reactions.^52^ This provides an experimental explanation for why ESPT in GFP is adiabatic.

**Figure 4.**
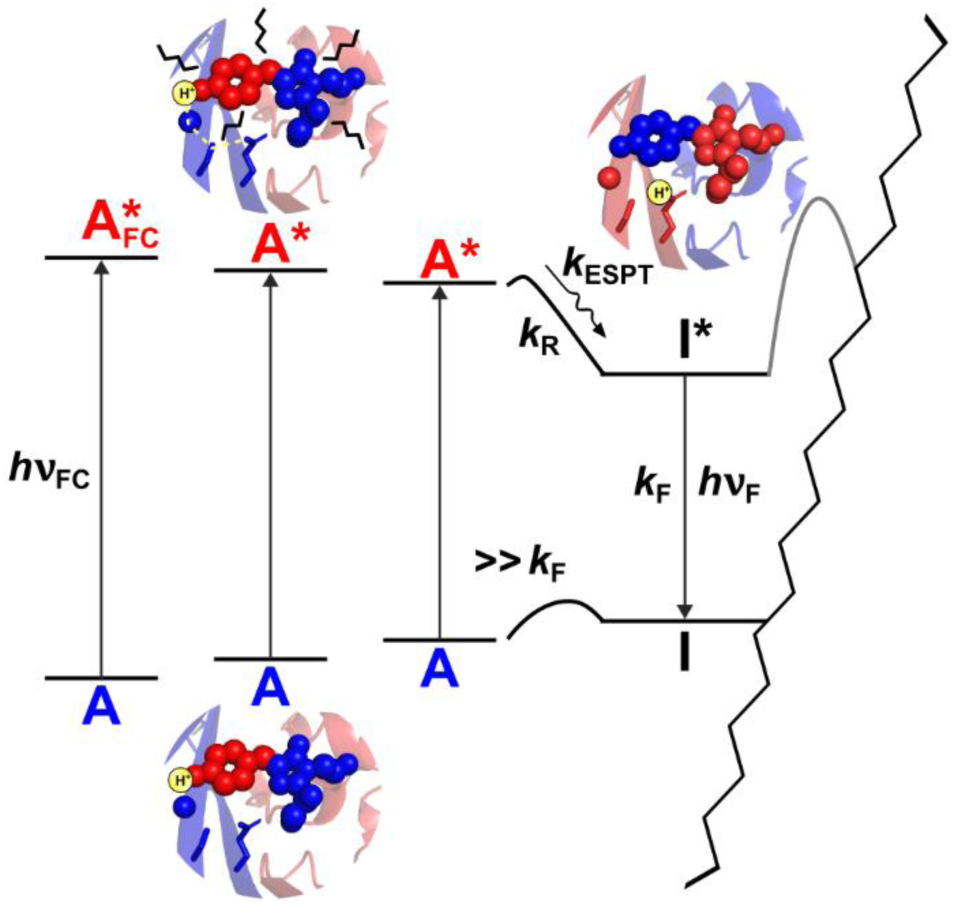
Different subpopulations of the neutral ground state (A, blue) with varying interaction energies between chromophore and environment dipole moments can, in principle, be excited by scanning across the *λ*_ex_ along the red edge. The rate of dipolar relaxation (*k*_R_) is, however, directly proportional to the sub-nanosecond rate of ESPT (*k*_ESPT_) between the neutral excited state (A*, red) and the emissive anionic excited state (I*, black). All emission (*k*_F_) therefore takes place from the relaxed, R state (*c.f.* Figure 1) regardless of *λ*_ex_, hence a flat REES profile. The B state side of the Jablonski diagram is cropped for clarity.

Upon excitation, the chromophore pKa is reduced by about 11.5 units.^35^ We therefore propose that rapid formation of this superacid chromophore forces the protein environment to respond, and could explain why the first proton transfer step in ESPT appears to be from S205 to E222 (i.e., in the environment) and not between the chromophore and the W22 water.^38, 54^ The superacid / dipolar relaxed state could correspond to the I_0_* state proposed by Di Donato et al.,^57^ which is proposed to contain a transient, low-barrier hydrogen bond (LBHB) within both the chromophore / water and the S205 / E222 donor-acceptor pairs of the proton wire.^56^ A LBHB places the transferring proton midway between donor and acceptor, with the energy minima of reactants and products being almost degenerate. This means the zero-point energy must be above the adiabatic electronic potential energy barrier and is thus likely to be sensitive to environmental dipolar effects that trigger the S205 to E222 proton transfer. The adiabatic ESPT we find in GFP is consistent with the reported existence of proton tunnelling,^54–55^ and could potentially help facilitate the exciton formation recently proposed between chromophores in GFP dimers.^58^

## Conclusions

There is mounting evidence that the asynchronous, concerted ESPT mechanism in GFP is very sensitive to slight modifications in its potential energy surface and requires vibrational relaxation and structural realignment to become favourable (alongside a tunnelling contribution).^54–56, 59–61^ Consistent with this sensitivity, for the first time we present experimental evidence that dipolar relaxation of the protein matrix during ESPT is also a key determinant of the reaction pathway and explains why it fits to an electronically adiabatic rate expression. This dipolar relaxation is likely to occur following vibrational relaxation and to be concomitant with structural realignment. Its adiabatic nature is consistent with the quantum dynamics proposed for the population of, and coupling between, the emissive states of GFP. There are also practical implications of this result for the design and use of light-activated proteins for bioimaging and optogenetics. In cases like GFP where the emission spectrum is essentially invariant irrespective of where along the red edge the chromophore is excited, the entire red edge could be excited at once using a broadband source, which would result in enhanced fluorescence at the same emission wavelengths. Moreover, understanding the intimate coupling between the protein matrix and an ultrafast chemical reaction provides a new perspective on how proteins can utilize environmental dynamics to guide reactions along a specific potential energy surface. This insight could inform the rational design of novel fluorescent proteins with tailored photophysics or the engineering of artificial enzymes or photoreceptors where controlling ultrafast charge transfer events is paramount.

## Experimental Section

See Supporting Information.

## ASSOCIATED CONTENT

## Supporting Information

Detailed experimental methods, and supporting figures.

## AUTHOR INFORMATION

## Notes

The authors declare no competing financial interest.

## ACKNOWLEDGMENT

ARJ thanks Prof. Steve Meech and Dr. Sussanah Borne-Worster for invaluable discussion and ARJ, MK, ISC and RWC thank Dr. Siddarth Shivkumar and Prof. Mike Shaw for feedback on the manuscript. ARJ thanks Steve Vogel and colleagues for providing the mAmetrine sample. ARJ, MK, ISC and RWC acknowledge the National Measurement System of the UK Government Department of Science, Innovation and Technology for funding. DDJ and AZ would like to thank the EPSRC (EP/V048147/1) for supporting this research and Cardiff School of Biosciences Protein Technology Hub for helping with the production of sfGFP H148C.

## ABBREVIATIONS

CSM: central spectral mass
CT: charge transfer
Em: emission
ESPT: excited state proton transfer
Ex: excitation
FC: Frank-Condon
GFP: green fluorescent protein
PT: proton transfer
R state: relaxed state
REES: red-edge excitation shift

## REFERENCES

1. Creed, D., The photophysics and photochemistry of the near-UV absorbing amino acids–I. tryptophan and its simple derivatives. Photochem. Photobiol. 1984, 39 (4), 537–562.

2. Camacho, I. S.; Theisen, A.; Johannissen, L. O.; Díaz-Ramos, L. A.; Christie, J. M.; Jenkins, G. I.; Bellina, B.; Barran, P.; Jones, A. R., Native mass spectrometry reveals the conformational diversity of the UVR8 photoreceptor. Proc. Natl. Acad. Sci. USA 2019, 116 (4), 1116–1125.

3. Rizzini, L.; Favory, J. J.; Cloix, C.; Faggionato, D.; O’Hara, A.; Kaiserli, E.; Baumeister, R.; Schafer, E.; Nagy, F.; Jenkins, G. I.; Ulm, R., Perception of UV-B by the *Arabidopsis* UVR8 protein. Science 2011, 332 (6025), 103–6.

4. Taniguchi, M.; Lindsey, J. S., Absorption and Fluorescence Spectral Database of Chlorophylls and Analogues. Photochem Photobiol 2021, 97 (1), 136–165.

5. Meech, S. R., Excited state reactions in fluorescent proteins. Chem Soc Rev 2009, 38 (10), 2922–2934.

6. van Thor, J. J.; Champion, P. M., Photoacid Dynamics in the Green Fluorescent Protein. Annu Rev Phys Chem 2023, 74 (Volume 74, 2023), 123–144.

7. Heijde, M.; Ulm, R., UV-B photoreceptor-mediated signalling in plants. Trends Plant Sci. 2012, 17 (4), 230–237.

8. Conrad, K. S.; Manahan, C. C.; Crane, B. R., Photochemistry of flavoprotein light sensors. Nat Chem Biol 2014, 10 (10), 801–809.

9. Jones, A. R., The photochemistry and photobiology of vitamin B_12_. Photochem. Photobiol. Sci. 2017, 16, 820–834.

10. Rockwell, N. C.; Lagarias, J. C., A Brief History of Phytochromes. ChemPhysChem 2010, 11 (6), 1172–1180.

11. Sies, H.; Mailloux, R. J.; Jakob, U., Fundamentals of redox regulation in biology. Nat Rev Mol Cell Biol 2024, 25 (9), 701–719.

12. Szczygiel, M.; Derewenda, U.; Scheiner, S.; Minor, W.; Derewenda, Z. S., A structural role for tryptophan in proteins, and the ubiquitous Trp C(δ1)-H…O=C (backbone) hydrogen bond. Acta crystallographica. Section D, Structural biology 2024, 80 (Pt 7), 551–562.

13. Lakowicz, J. R., Mechanisms and Dynamics of Solvent Relaxation. In Principles of Fluorescence Spectroscopy, Springer US: Boston, MA, 1983; pp 217–255.

14. Demchenko, A. P., Weber’s Red-Edge Effect that Changed the Paradigm in Photophysics and Photochemistry. In Perspectives on Fluorescence: A Tribute to Gregorio Weber, Jameson, D. M., Ed. Springer International Publishing: Cham, 2016; pp 95–141.

15. Demchenko, A. P., The red-edge effects: 30 years of exploration. Luminescence 2002, 17 (1), 19–42.

16. Chattopadhyay, A.; Haldar, S., Dynamic Insight into Protein Structure Utilizing Red Edge Excitation Shift. Acc. Chem. Res. 2014, 47 (1), 12–19.

17. Demchenko, A. P.; Ladokhin, A. S., Red-edge-excitation fluorescence spectroscopy of indole and tryptophan. Eur Biophys J 1988, 15 (6), 369–379.

18. Demchenko, A. P., Red-edge-excitation fluorescence spectroscopy of single-tryptophan proteins. Eur Biophys J 1988, 16 (2), 121–129.

19. Mukherjee, S.; Chattopadhyay, A., Wavelength-selective fluorescence as a novel tool to study organization and dynamics in complex biological systems. J. Fluoresc. 1995, 5 (3), 237–246.

20. Kwok, A.; Camacho, I.; Winter, S.; Knight, M.; Meade, R.; Van der Kamp, M.; Turner, A.; O’Hara, J.; Mason, J.; Jones, A.; Arcus, V.; Pudney, C., A Thermodynamic Model for Interpreting Tryptophan Excitation-Energy-Dependent Fluorescence Spectra Provides Insight Into Protein Conformational Sampling and Stability. Front. Mol. Biosci. 2021, 8, 778244.

21. Warrender, A. K.; Pan, J.; Pudney, C.; Arcus, V. L.; Kelton, W., Red edge excitation shift spectroscopy is highly sensitive to tryptophan composition. J. R. Soc. Interface 2023, 20 (208), 20230337.

22. Ramakrishnan, K.; Johnson, R. L.; Winter, S. D.; Worthy, H. L.; Thomas, C.; Humer, D. C.; Spadiut, O.; Hindson, S. H.; Wells, S.; Barratt, A. H.; Menzies, G. E.; Pudney, C. R.; Jones, D. D., Glycosylation increases active site rigidity leading to improved enzyme stability and turnover. FEBS J 2023, 290 (15), 3812–3827.

23. Remington, S. J., Fluorescent proteins: maturation, photochemistry and photophysics. Curr. Opin. Struct. Biol. 2006, 16 (6), 714–721.

24. van Thor, J. J., Photoreactions and dynamics of the green fluorescent protein. Chem. Soc. Rev. 2009, 38 (10), 2935–2950.

25. Acharya, A.; Bogdanov, A. M.; Grigorenko, B. L.; Bravaya, K. B.; Nemukhin, A. V.; Lukyanov, K. A.; Krylov, A. I., Photoinduced Chemistry in Fluorescent Proteins: Curse or Blessing? Chem. Rev. 2017, 117 (2), 758–795.

26. Heim, R.; Prasher, D. C.; Tsien, R. Y., Wavelength mutations and posttranslational autoxidation of green fluorescent protein. Proc Nat Acad Sci USA 1994, 91 (26), 12501–12504.

27. Yang, F.; Moss, L. G.; Phillips, G. N., The molecular structure of green fluorescent protein. Nat Biotech 1996, 14 (10), 1246–1251.

28. Morise, H.; Shimomura, O.; Johnson, F. H.; Winant, J., Intermolecular energy transfer in the bioluminescent system of Aequorea. Biochemistry 1974, 13 (12), 2656–2662.

29. Brejc, K.; Sixma, T. K.; Kitts, P. A.; Kain, S. R.; Tsien, R. Y.; Ormö, M.; Remington, S. J., Structural basis for dual excitation and photoisomerization of the *Aequorea victoria* green fluorescent protein. Proc Nat Acad Sci USA 1997, 94 (6), 2306–2311.

30. Hartley, A. M.; Worthy, H. L.; Reddington, S. C.; Rizkallah, P. J.; Jones, D. D., Molecular basis for functional switching of GFP by two disparate non-native post-translational modifications of a phenyl azide reaction handle. Chem. Sci. 2016, 7 (10), 6484–6491.

31. Worthy, H. L.; Auhim, H. S.; Jamieson, W. D.; Pope, J. R.; Wall, A.; Batchelor, R.; Johnson, R. L.; Watkins, D. W.; Rizkallah, P.; Castell, O. K.; Jones, D. D., Positive functional synergy of structurally integrated artificial protein dimers assembled by Click chemistry. Commun. Chem. 2019, 2 (1), 83.

32. Ahmed, R. D.; Jamieson, W. D.; Vitsupakorn, D.; Zitti, A.; Pawson, K. A.; Castell, O. K.; Watson, P. D.; Jones, D. D., Molecular dynamics guided identification of a brighter variant of superfolder Green Fluorescent Protein with increased photobleaching resistance. Commun. Chem. 2025, 8 (1), 174.

33. Elsliger, M.-A.; Wachter, R. M.; Hanson, G. T.; Kallio, K.; Remington, S. J., Structural and Spectral Response of Green Fluorescent Protein Variants to Changes in pH. Biochemistry 1999, 38 (17), 5296–5301.

34. Tsien, R. Y., The green fluorescent protein. Annu Rev Biochem 1998, 67.

35. Chattoraj, M.; King, B. A.; Bublitz, G. U.; Boxer, S. G., Ultra-fast excited state dynamics in green fluorescent protein: multiple states and proton transfer. Proc Nat Acad Sci USA 1996, 93 (16), 8362–8367.

36. Creemers, T. M. H.; Lock, A. J.; Subramaniam, V.; Jovin, T. M.; Völker, S., Three photoconvertible forms of green fluorescent protein identified by spectral hole-burning. Nat Struct Biol 1999, 6 (6), 557–560.

37. Palm, G. J.; Zdanov, A.; Gaitanaris, G. A.; Stauber, R.; Pavlakis, G. N.; Wlodawer, A., The structural basis for spectral variations in green fluorescent protein. Nat Struct Biol 1997, 4 (5), 361–365.

38. Lill, M. A.; Helms, V., Proton shuttle in green fluorescent protein studied by dynamic simulations. Proc Nat Acad Sci USA 2002, 99 (5), 2778–2781.

39. Pédelacq, J.-D.; Cabantous, S.; Tran, T.; Terwilliger, T. C.; Waldo, G. S., Engineering and characterization of a superfolder green fluorescent protein. Nat. Biotech. 2006, 24 (1), 79–88.

40. Ahmed, R. D.; Vitsupakorn, D.; Hartwell, K. D.; Albalawi, K.; Rizkallah, P.; Watson, P. D.; Jones, D. D., Chromophore charge-state switching through copper-dependent homodimerisation of an engineered green fluorescent protein. bioRxiv 2025, 2025.08.27.672602.

41. Ai, H.-w.; Hazelwood, K. L.; Davidson, M. W.; Campbell, R. E., Fluorescent protein FRET pairs for ratiometric imaging of dual biosensors. Nat. Meth. 2008, 5 (5), 401–403.

42. Lakowicz, J. R.; Keating-Nakamoto, S., Red-edge excitation of fluorescence and dynamic properties of proteins and membranes. Biochemistry 1984, 23 (13), 3013–3021.

43. Mandal, D.; Tahara, T.; Meech, S. R., Excited-State Dynamics in the Green Fluorescent Protein Chromophore. J. Phys. Chem. B 2004, 108 (3), 1102–1108.

44. Volkmer, A.; Subramaniam, V.; Birch, D. J. S.; Jovin, T. M., One- and Two-Photon Excited Fluorescence Lifetimes and Anisotropy Decays of Green Fluorescent Proteins. Biophys. J. 2000, 78 (3), 1589–1598.

45. Pan, C.-P.; Muiño, P. L.; Barkley, M. D.; Callis, P. R., Correlation of Tryptophan Fluorescence Spectral Shifts and Lifetimes Arising Directly from Heterogeneous Environment. J. Phys. Chem. B 2011, 115 (12), 3245–3253.

46. Haldar, S.; Chattopadhyay, A., Dipolar Relaxation within the Protein Matrix of the Green Fluorescent Protein: A Red Edge Excitation Shift Study. J. Phys. Chem. B 2007, 111 (51), 14436–14439.

47. Nifosì, R.; Tozzini, V., Molecular dynamics simulations of enhanced green fluorescent proteins: Effects of F64L, S65T and T203Y mutations on the ground-state proton equilibria. Proteins 2003, 51 (3), 378–389.

48. Arpino, J. A. J.; Rizkallah, P. J.; Jones, D. D., Crystal Structure of Enhanced Green Fluorescent Protein to 1.35 Å Resolution Reveals Alternative Conformations for Glu222. PLOS ONE 2012, 7 (10), e47132.

49. Demchenko, A. P.; Sytnik, A. I., Solvent reorganizational red-edge effect in intramolecular electron transfer. Proc Nat Acad Sci USA 1991, 88 (20), 9311–9314.

50. Demchenko, A. P.; Sytnik, A. I., Site selectivity in excited-state reactions in solutions. J. Phys. Chem. 1991, 95 (25), 10518–10524.

51. Demchenko, A. P., Protein fluorescence, dynamics and function: exploration of analogy between electronically excited and biocatalytic transition states. Biochim. Biophys. Acta 1994, 1209 (2), 149–164.

52. Maroncelli, M.; MacInnis, J.; Fleming, G. R., Polar Solvent Dynamics and Electron-Transfer Reactions. Science 1989, 243 (4899), 1674–1681.

53. Kennis, J. T. M.; Larsen, D. S.; van Stokkum, I. H. M.; Vengris, M.; van Thor, J. J.; van Grondelle, R., Uncovering the hidden ground state of green fluorescent protein. Proc Nat Acad Sci USA 2004, 101 (52), 17988–17993.

54. Bourne-Worster, S.; Worth, G. A., Quantum dynamics of excited state proton transfer in green fluorescent protein. J. Chem. Phys. 2024, 160 (6).

55. Salna, B.; Benabbas, A.; Sage, J. T.; van Thor, J.; Champion, P. M., Wide-dynamic-range kinetic investigations of deep proton tunnelling in proteins. Nat. Chem. 2016, 8 (9), 874–880.

56. Nadal-Ferret, M.; Gelabert, R.; Moreno, M.; Lluch, J. M., Transient low-barrier hydrogen bond in the photoactive state of green fluorescent protein. Phys. Chem. Chem. Phys. 2015, 17 (46), 30876–30888.

57. Di Donato, M.; van Wilderen, L. J. G. W.; Van Stokkum, I. H. M.; Stuart, T. C.; Kennis, J. T. M.; Hellingwerf, K. J.; van Grondelle, R.; Groot, M. L., Proton transfer events in GFP. Phys. Chem. Chem. Phys. 2011, 13 (36), 16295–16305.

58. Nguyen, T.; Puhl, H. L.; Chen, E.; Hines, K.; Rani, C.; Blank, P. S.; Carlotti, B.; Kim, Y.; Goodson, T.; Vogel, S. S., Anomalous photophysical behaviors attributed to excitonic coupling in fluorescent proteins. Biophys. J. 2025.

59. Petrone, A.; Cimino, P.; Donati, G.; Hratchian, H. P.; Frisch, M. J.; Rega, N., On the Driving Force of the Excited-State Proton Shuttle in the Green Fluorescent Protein: A Time-Dependent Density Functional Theory (TD-DFT) Study of the Intrinsic Reaction Path. J. Chem. Theory Comput. 2016, 12 (10), 4925–4933.

60. Donati, G.; Petrone, A.; Caruso, P.; Rega, N., The mechanism of a green fluorescent protein proton shuttle unveiled in the time-resolved frequency domain by excited state ab initio dynamics. Chem. Sci. 2018, 9 (5), 1126–1135.

61. Fang, C.; Frontiera, R. R.; Tran, R.; Mathies, R. A., Mapping GFP structure evolution during proton transfer with femtosecond Raman spectroscopy. Nature 2009, 462 (7270), 200–204.

