## Supporting Information for "Environmental dipolar relaxation during excited state proton transfer in Green Fluorescent Protein"

EXPERIMENTAL METHODS

**Materials**

L-tryptophan was purchased from Formedium and wild-type *Av*GFP from Invitrogen; both were used without further purification. Trp was dissolved into PBS buffer (Fisher BioReagents), filtered with 0.22 µm syringe filter (Sartorius Minisart™). *Av*GFP was exchanged into 50 mM Tris-HCl, pH = 8 buffer by diluting it into the buffer and then concentrating it with Vivaspin 20 centrifugal concentrator (10 kDa cutoff) for 5 min at 4000 xg. The cycle was repeated three more times. After the buffer exchange, the sample was filtered with 0.22 μm Ultrafree centrifugal filter (Millipore) and diluted to a suitable concentration for the UV-Vis and fluorescence measurements.

**Protein expression and purification**

The sfGFP H148C variant[1] (based on the pBAD vector) was generated *via* site-directed mutagenesis using a whole plasmid inverse PCR process as described previously.[2-3] Polymerase chain reaction (PCR) was run following the Q5® High Fidelity DNA Polymerase (NEB #M0491) protocol. Recombinant production was performed in chemically competent *E. coli* TOP10™ cells, which were transformed with the plasmid and plated on LB agar plates supplemented with 100 μg/mL ampicillin. A single colony was taken and used to inoculate a 10 mL 2xYT starter culture supplemented with antibiotic and incubated overnight with shaking. The overnight culture was used to inoculate a 1L culture of autoinduction media (0.5% (v/v) glycerol; 0.05% (w/v) glucose; 0.2% lactose; 25 mM $Na_2HPO_4$; 25 mM $KH_2PO_4$; 50 mM $NH_4Cl$; 5 mM $NaSO_4$; 2 mM $MgSO_4$; 0.05% (w/v) L-arabinose). The culture was inoculated overnight at 37 °C with shaking. Cells were harvested by centrifugation at 5000 xg for 20 minutes at 4 °C and suspended in 20 mL 50 mM Tris, pH = 8.0. Cells were lysed using the French Pressure Cell. Soluble cell lysate was separated from insoluble fractions *via* centrifugation at 25000 xg for 1 hour. Protein purification was carried out with an ÄKTA Purifier FPLC using columns purchased from Cytiva and protein elution monitored at 280 nm, 400 nm and 485 nm. Clarified cell lysate was first passed through a 5 mL His Trap™ HP column equilibrated in 50 mM Tris, pH = 8.0 buffer containing 10 mM Imidazole. Bound target protein was then eluted by the addition of the elution buffer containing imidazole at a gradient from 10 to 500 mM imidazole. Fractions were then checked for purity via SDS-PAGE analysis. Finally, the proteins were further purified by size exclusion chromatography (SEC) using a HiLoad™ 16/600 Superdex™ S75 pg column (Cytiva) equilibrated with 50 mM Tris, pH = 8.0. protein samples were concentrated using Amicon® 30 kDa Ultra Centrifugal Filter (Merck, Millipore) by centrifugation at 3500 xg until the desired volume is reached.

mAmetrine[4] protein samples were gifted by the Vogel Lab at NIH. All GFP samples were diluted in 50 mM Tris-HCl, pH = 8, prior to use.

**Spectroscopy**

Absorption spectra of Trp and GFP variants were measured using a Cary 60 UV-vis Spectrophotometer (Agilent) in 10 mm Quartz cuvette (Starna) with corresponding buffer background and 600 nm/min scanning speed.

Most of the fluorescence REES data were measured using a FLS1000 Photoluminescence Spectrometer (Edinburgh Instruments) using a 10 mm path length Quartz cuvette (Starna), although sfGFP H148C data were acquired using a LS55 Fluorescence Spectrometer (PerkinElmer). For all GFP variants, the measurement parameters were: 5 nm excitation and emission slit widths, 0.5 nm steps in emission wavelength, 0.12 s/nm integration time. Spectra were measured at +20 °C and +50 °C (after > 10 min incubation at the set temperature), maintained using a temperature-controlled cuvette holder connected to a water bath, to study the temperature dependence of the REES profile in GFP.[5] *Av*GFP and sfGFP H148C were excited from 400 to 440 nm by 2 nm steps, and emission collected from 495 to 630 nm. mAmetrine was excited from 404 to 460 nm by 2 nm steps, and emission was collected from 490 to 670 nm. For Figure S3, *Av*GFP was excited from 410 nm to 480 nm by 5 nm steps, and emission was collected from 490 to 650 nm, and only a single dataset was collected at +20 °C.

Trp was diluted to a final concentration of 266 μM resulting in absorbance of 1 at 290 nm. This concentration enabled the acquisition of a non-negligible fluorescence signal at the longest excitation wavelengths. The REES data were measured with FLS1000 at +20 °C by exciting from 290 nm to 330 nm and collecting emission from 340 to 550 nm, both with 1 nm steps and 0.25 s/nm integration time. The excitation and emission slit widths were set to 2.20 nm and 1.00 nm, respectively.

All REES data were collected in triplicates. For all samples, an identical dataset of buffer alone was measured in triplicate to allow for buffer correction.

**REES Data Analysis**

Analysis of the REES data was performed using MATLAB (R2024b) as described previously.[6] To correct for the water Raman peak, the corresponding buffer spectrum was subtracted from each emission spectrum. Then for each buffer-corrected spectrum the Centre of Spectral Mass (CSM) was calculated according to Eq 1

$$CSM = \frac{\Sigma(f_i \times \lambda_{Em})}{\Sigma f_i} \qquad (1)$$

For each excitation wavelength the mean and standard error of the CSM was then calculated from triplicates of such values.

**Electronic Dipole Calculations**

Chromophores of *Av*GFP and sfGFP were extracted from available crystal structures (PDB entries 1GFL and 2B3P, respectively) and geometry optimised without symmetry constraints at PBE0/def2-TZVP level.[7-8] The CRO environment was modelled as point charges of nearby residues and water molecules, inside the CPCM model of aqueous solvation.[9] Included residues were Arg96, His148, Ser205 and Glu222, along with waters 248, 259, 262 and 264 (for *Av*GFP) and waters 815, 821, 828 and 843 (for sfGFP). Absorption spectra and ground/excited state dipole moment information at the same level, using the relaxed density for the excited state and non-equilibrium solvation. Trp data were calculated for the isolated, amide-capped amino acid only. All calculations were performed in Orca 6.1.0.[10]

The figures illustrating the dipoles of the chromophores in the different GFP variants were created using the relevant PDB file (see above) as a starting point. Avogadro[11] was used to add the molecules of water and VMD[12] to export the file into a .obj file. Blender[13] was used to read the file and the vector arrows were added using a custom Python software to create the vector composition. The orientations and magnitude of each dipole are based in quantum mechanical calculations described above.

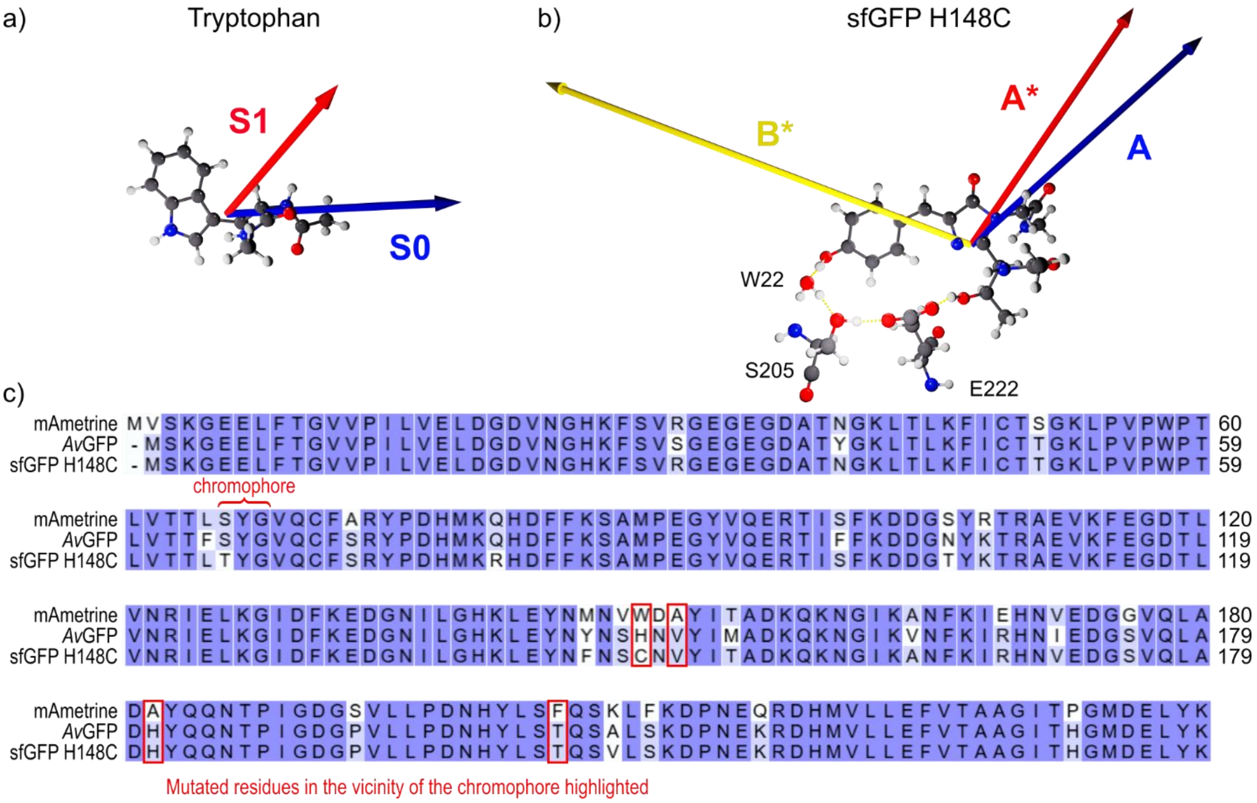


**Figure S1. Calculated electronic dipoles** of a) tryptophan in ground state (S0, blue) and excited state (S1, red); and b) sfGFP H148C for the neutral ground (A, blue) and excited (A*, red) states, and anionic excited state (B*, yellow). The sfGFP H148C chromophore (T65-Y66-G67) is shown in the local environment with a water molecule (W22), and two residues, S205 and E222, involved in the ESPT, highlighted with dashed lines. c) **Sequence alignment** of the three GFP variants tested: mAmetrine, *Av*GFP, and sfGFP H148C. The mutated residues whose side chains are in proximity of the chromophore based on the available crystal structures are highlighted in red.

**Table S1. Calculated transition dipole moments** (magnitude and direction) of the ground and different excited states of tryptophan (Trp) and GFP variants *Av*GFP, and sfGFP H148C. The dipole moment for mAmetrine was not calculated due to the lack of a crystal structure with which to accurately model the environment.

| Chromophore | State | Dipole (a.u.) | x | y | z | Θ (°) |
|---|---|---|---|---|---|---|
| **Trp** | G | 2.79 | -2.64 | -0.34 | 0.84 | |
| | S1 | 2.23 | -1.14 | 0.49 | 1.85 | 45.1 |
| ***Av*GFP** | A | 5.08 | -3.46 | 3.65 | 0.72 | |
| | A* | 5.93 | -4.36 | 3.93 | 0.85 | 4.5 |
| | B | 7.78 | 7.53 | 1.98 | -0.07 | |
| | B* | 6.07 | 5.71 | 2.05 | 0.00 | 5.1 |
| **sfGFP H148C** | A | 4.47 | -3.30 | 2.68 | 1.38 | |
| | A* | 4.16 | -2.33 | 3.06 | 1.59 | 13.5 |
| | B | 8.53 | 8.23 | 1.99 | 0.98 | |
| | B* | 6.86 | 6.42 | 2.16 | 1.11 | 5.6 |

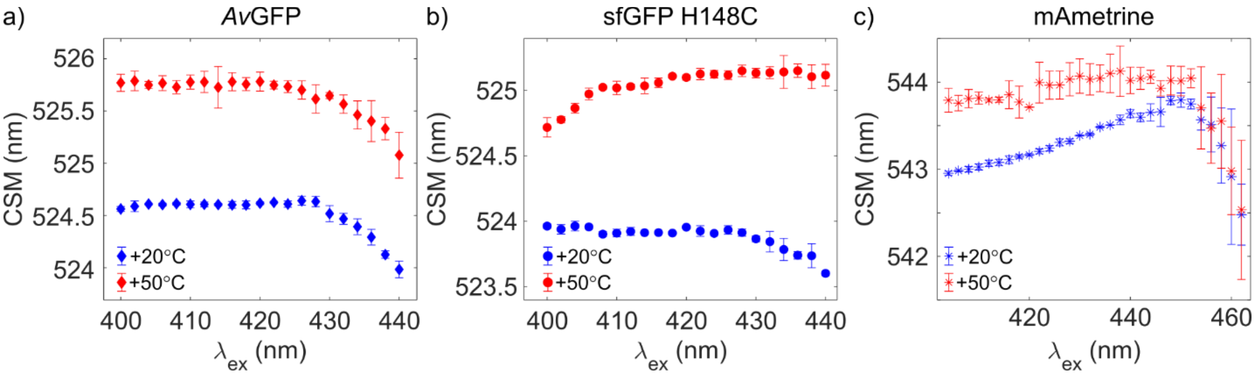


**Figure S2. Zoomed-in CSM plots** of a) *Av*GFP, b) sfGFP H148C, and c) mAmetrine in Figure 3b in the main paper at +20 °C (blue) and at +50 °C (red).

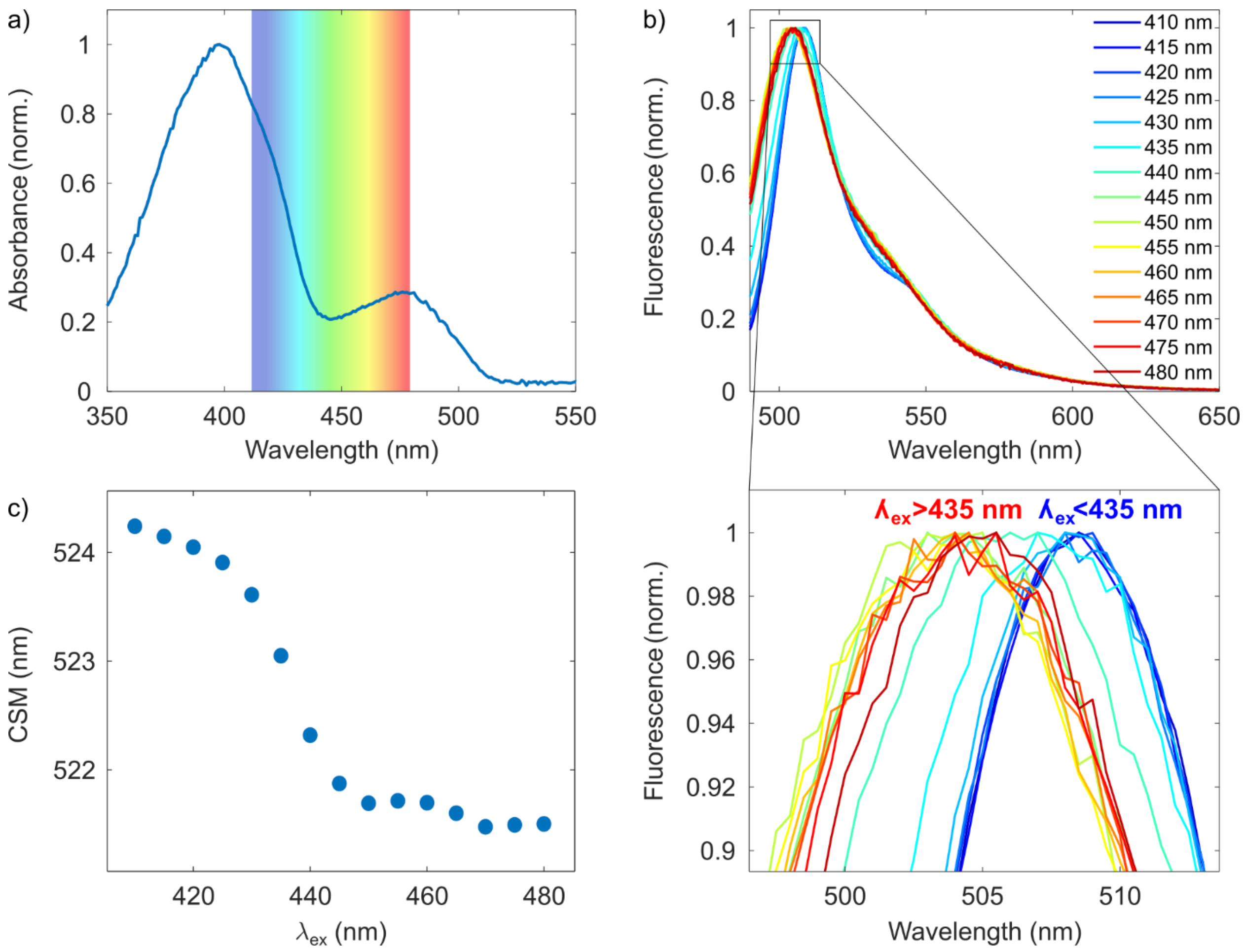


**Figure S3. Extended excitation along the red edge of the A band of *Av*GFP into the B band.** a) Absorption spectrum of *Av*GFP at +20 °C. The rainbow block highlights the excitation wavelengths used to collect REES data. b) The normalised emission spectra of *Av*GFP from 410 to 480 nm. The zoomed-in figure in the bottom shows the emission peak maxima and the blue shift from 509 nm to 503-504 nm when moving to excitation wavelengths above 435 nm, corresponding to the second absorption peak. c) CSM of the *Av*GFP emission spectra plot as a function of $\lambda_{ex}$, which demonstrates the CSM shift taking place between 430 and 440 nm.

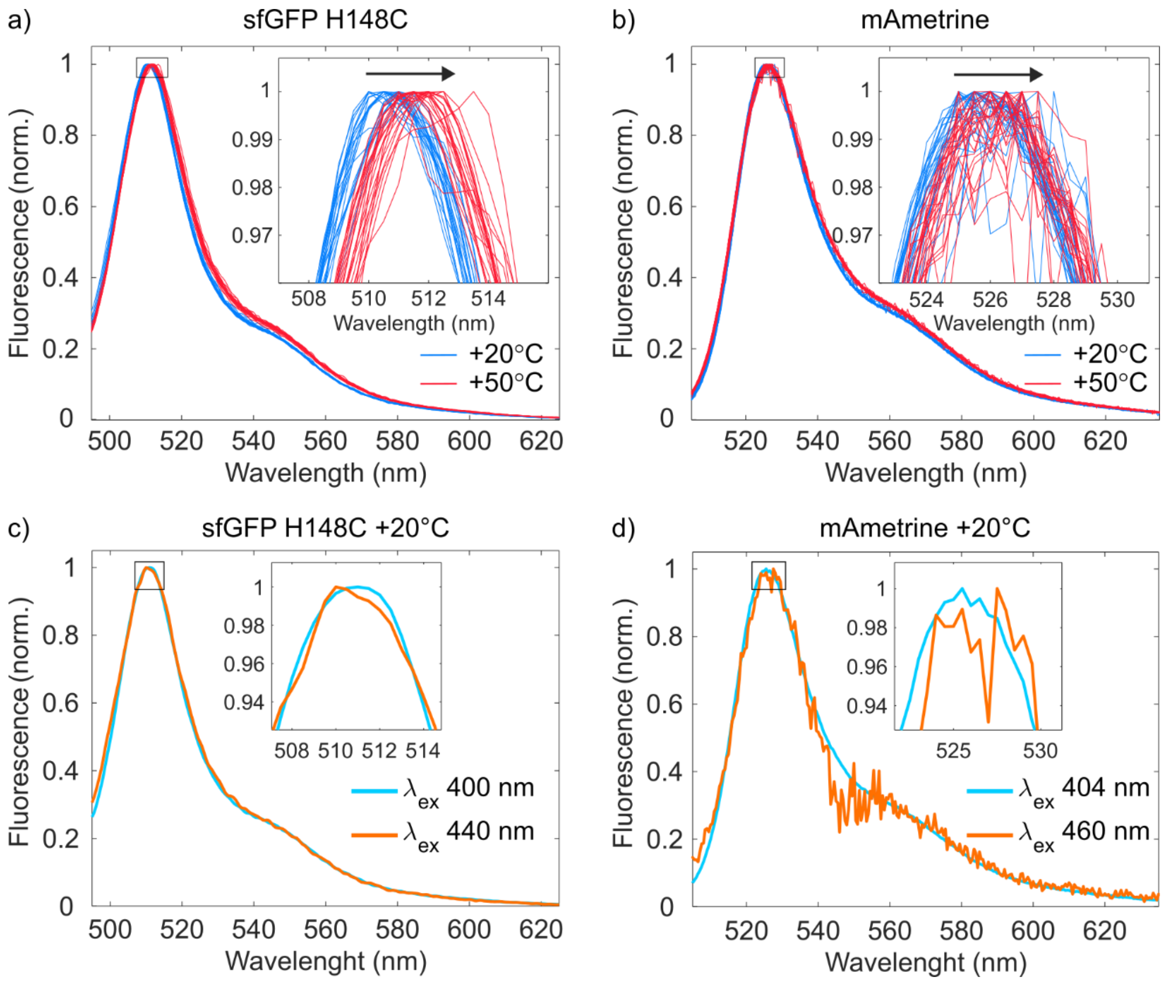


**Figure S4. Emission spectra** of a) sfGFP H148C with excitation wavelengths from 400 to 440 nm with 2 nm steps, and b) mAmetrine at excitation wavelengths from 404 to 450 nm with 2 nm steps, both at +20 °C (blue) and +50 °C (red). c-d) Overlaid emission spectra from of the shortest (cyan) and longest (orange) excitation wavelength in the REES matrix for c) sfGFP H148C and d) mAmetrine at +20 °C with peak position enlarged (inset). In both, the longest excitation broadens the spectrum in the blue side of the peak. This explains the ultimate blue-shift of the CSM as a function of $\lambda_{ex}$ (Figure S2b&c) as emission from the B* state begins to contribute to the signal, despite the lack of a blue-shifted peak wavelength. For mAmetrine, there is evidence of non-neglible signal from the water Raman peak, despite buffer correction. This may distort the magnitude of the CSM blue shift in Figure S2c.

REFERENCES


1. Ahmed, R. D.; Vitsupakorn, D.; Hartwell, K. D.; Albalawi, K.; Rizkallah, P.; Watson, P. D.; Jones, D. D., Chromophore charge-state switching through copper-dependent homodimerisation of an engineered green fluorescent protein. *bioRxiv* **2025**, 2025.08.27.672602.
2. Reddington, S. C.; Rizkallah, P. J.; Watson, P. D.; Pearson, R.; Tippmann, E. M.; Jones, D. D., Different Photochemical Events of a Genetically Encoded Phenyl Azide Define and Modulate GFP Fluorescence. *Angew Chem Int Ed* **2013,** *52* (23), 5974-5977.
3. Worthy, H. L.; Auhim, H. S.; Jamieson, W. D.; Pope, J. R.; Wall, A.; Batchelor, R.; Johnson, R. L.; Watkins, D. W.; Rizkallah, P.; Castell, O. K.; Jones, D. D., Positive functional synergy of structurally integrated artificial protein dimers assembled by Click chemistry. *Commun. Chem.* **2019,** *2* (1), 83.
4. Ai, H.-w.; Hazelwood, K. L.; Davidson, M. W.; Campbell, R. E., Fluorescent protein FRET pairs for ratiometric imaging of dual biosensors. *Nat. Meth.* **2008,** *5* (5), 401-403.
5. Haldar, S.; Chattopadhyay, A., Dipolar Relaxation within the Protein Matrix of the Green Fluorescent Protein: A Red Edge Excitation Shift Study. *J. Phys. Chem. B* **2007,** *111* (51), 14436-14439.
6. Knight, M. J.; Woolley, R. E.; Kwok, A.; Parsons, S.; Jones, H. B. L.; Gulácsy, C. E.; Phaal, P.; Kassaar, O.; Dawkins, K.; Rodriguez, E.; Marques, A.; Bowsher, L.; Wells, S. A.; Watts, A.; van den Elsen, J. M. H.; Turner, A.; O'Hara, J.; Pudney, C. R., Monoclonal antibody stability can be usefully monitored using the excitation-energy-dependent fluorescence edge-shift. *Biochem. J.* **2020,** *477* (18), 3599-3612.
7. Adamo, C.; Barone, V., Toward reliable density functional methods without adjustable parameters: The PBE0 model. *J. Chem. Phys.* **1999,** *110* (13), 6158-6170.
8. Weigend, F.; Ahlrichs, R., Balanced basis sets of split valence, triple zeta valence and quadruple zeta valence quality for H to Rn: Design and assessment of accuracy. *Phys Chem Chem Phys* **2005,** *7* (18), 3297-3305.
9. Cammi, R.; Mennucci, B.; Tomasi, J., Fast Evaluation of Geometries and Properties of Excited Molecules in Solution: A Tamm-Dancoff Model with Application to 4-Dimethylaminobenzonitrile. *J Phys Chem A* **2000,** *104* (23), 5631-5637.
10. Neese, F., Software Update: The ORCA Program System—Version 6.0. *Comput Mol Sci* **2025,** *15* (2), e70019.
11. Hanwell, M. D.; Curtis, D. E.; Lonie, D. C.; Vandermeersch, T.; Zurek, E.; Hutchison, G. R., Avogadro: an advanced semantic chemical editor, visualization, and analysis platform. *J. Cheminform.* **2012,** *4* (1), 17.
12. Humphrey, W.; Dalke, A.; Schulten, K., VMD: Visual molecular dynamics. *J. Mol. Graphics* **1996,** *14* (1), 33-38.
13. Community, B. O. *Blender - a 3D modelling and rendering package*, 3.2; Stichting Blender Foundation: Amsterdam, 2018.
